# Prolonged over-expression of PLK4 amplifies centrosomes through formation of inter-connected centrosome rosette clusters

**DOI:** 10.1101/2023.10.10.561779

**Authors:** Selahattin Can Özcan, Batuhan Mert Kalkan, Enes Çiçek, Ata Alpay Canbaz, Ceyda Açılan Ayhan

## Abstract

The centrosome cycle is a tightly regulated process to ensure proper segregation of chromosomes. Not surprisingly, centriole number is tightly controlled via multiple mechanisms, one of which involves PLK4, an upstream kinase facilitating centriole biogenesis and duplication. Aberrations in this process can result in supernumerary centrosomes, which are frequently observed in a variety of cancers due to high levels of PLK4. Interestingly, extra centrosomes induced by PLK4 over-expression go through unique intermediate structures called the centrosome rosettes (CRs), where the mother centriole is surrounded by numerous daughter centrioles. The maturation and molecular nature of these CRs have not been investigated in detail. Upon prolonged PLK4 over-expression, cells exhibited large centrosomes that were clustered and contained more than two CRs, which we defined as centrosome rosette clusters (CRCs). As expected, these structures required high PLK4 levels at two consecutive cell cycles and were still interconnected with canonical centrosomal linker proteins such as C-Nap1, Rootletin, and Cep68. Knockout of these linker proteins resulted in distancing of CRs and CRCs as observed by increased diameter of the CRCs in interphase. In contrast, Nek2 knockout inhibited the separation of CRCs in prometaphase, providing functional evidence for the binding of CRC structures with centrosomal linker proteins. These results suggest a cell cycle dependent model for PLK4 induced centrosome amplification, which occurs in two consecutive cell cycles: (i) CR state in the first cell cycle, and (ii) CRC state in the second cell cycle.

**Author summary:** The overexpression of PLK4 can lead to the formation of centrosome rosette structures, which harbor two centrioles around the mother centriole. Although the generation of centrosome rosettes by PLK4 overexpression has been previously investigated, little is known about the cell cycle-dependent maturation and linking of these structures. Here, we report that prolonged PLK4 overexpression results in amplification of centrosomes through the generation of centrosome rosette clusters (CRCs). These CRCs are interconnected via canonical centrosomal linker proteins such as C-Nap1, Rootletin, and CEP68 and are regulated by mechanisms controlling centrosome linking and separation. We also describe two different spatial binding types of amplified centrosomes following PLK4 induction: planar-oriented and circular-oriented. Since PLK4-associated centrosome amplification occurs naturally in both cancer and multiciliated cells, we believe that this research will contribute to a better understanding of the canonical mechanism of PLK4-induced centrosome amplification.

## Introduction

Centrosomes, composed of two centrioles and pericentriolar material (PCM), are the primary microtubule organizing centers in animal cells [1]. Centrosomes play essential roles in coordinating interphase microtubule organization and forming spindle poles during mitosis, ensuring proper chromosome segregation [2]. Additionally, centrosomes serve as nucleating basal bodies for ciliagenesis in non-proliferating cells. The mature-mother centriole, one of the two centrioles in a G1 cell, is decorated with subdistal and distal appendages, enabling the generation of cilia by docking to the plasma membrane [3]. Cilia are vital for cell signaling, fluid movement, and sensory perception, and defects in their formation or function can lead to various human diseases known as ciliopathies [4]. Moreover, centrosomes are involved in cell polarity and migration, and abnormalities in these processes can contribute to cancer and other diseases [5].

Centriole and centrosome numbers in a cell are tightly controlled, with centrosomes duplicating once per cell cycle and newly formed centrioles maturing as the cell progresses through the cycle. The centrosome cycle in animal cells consists of five consecutive events, known as (i) centriole biogenesis, (ii) elongation, (iii) separation, (iv) disengagement, and (v) maturation [6]. Briefly, recently divided G1 cells contain two centrioles in different stages of maturation: the younger mother, which was assembled in the previous cell cycle, and the mature mother, which was assembled one cycle earlier. These two mother centrioles are linked by a proteinaceous linker, called the centrosomal linker [7]. At the G1-S transition, procentrioles start to grow on a single site at the proximal part of each mother centriole. Short procentrioles become daughter centrioles through an elongation process that proceeds until G2. Mother centrioles are separated during prophase by phosphorylation of centrosomal linker proteins and form spindle poles in mitosis. In late mitosis, daughter centrioles become disengaged from mothers, and both centrioles of a centrosome mature to form two mother (one mature and one young) centrioles of the daughter cells.

Centrosome abnormalities are a common occurrence in cancer, with the most well-documented abnormality being centrosome amplification. Centrosome amplification is one of the major causes of chromosome missegregation and chromosomal instability in cancers [8]. It can arise from various mechanisms, including (i) activation of oncogenes [9]; (ii) loss of tumor suppressor factors [10]; (iii) cell division errors that can result from defects in spindle assembly checkpoint function, cytokinesis failure, or other mitotic defects [11]; and (iv) hypoxic conditions in the tumor microenvironment, which can disrupt normal centrosome function and promote centrosome amplification [12]. Thus, centrosome amplification is a multifactorial process that contributes to the development and progression of various cancers and has the potential to be a promising target for therapeutic intervention.

PLK4 is considered as the master regulator of centriole duplication and is responsible for initiating the formation of a procentriole on the mother centriole wall. Studies on the role of PLK4 in centriole biogenesis have mostly been conducted by its over-expression, resulting in the formation of centrosome rosettes, which are a flower-shaped cluster of procentrioles on the mother centriole wall [13–15]. The first report of endogenous formation of centrosome rosettes (CRs) in ciliated mouse oviducts dates back to 1971 [14]. Subsequent studies showed the presence of centrosome rosettes in primary tumor samples of multiple myeloma, glioblastoma, and colon cancers [16]. Endogenous centrosome rosette formation has also been observed in olfactory sensory neuron precursor cells [17]. These findings clearly demonstrate that PLK4-induced CRs are readily detectable in both cancer and stem cells, implying its importance for cell biology.

Previous research has successfully characterized the generation of centrosome rosettes through the over-expression of PLK4 or STIL. However, the maturation and cell cycle-dependent cycling of CR structures have remained unexplained. To better understand the cell cycle-dependent CR biology, we characterized the structure and linking of PLK4-induced CRs using high-resolution confocal microscopy. Our observations revealed that long-term PLK4 induction generates cells with more than two centrosome rosettes linked to each other with centrosomal linkers, which we named as centrosome rosette clusters (CRCs). We then investigated the cell cycle-dependent maturation of CRs into CRCs and found that CRCs behave similarly to normal centrosomes in terms of maturation, cohesion, and separation. Furthermore, we demonstrated the functional necessity of centrosomal linkers for bridging CRCs in CRISPR/Cas9-guided C-Nap1, Rootletin, and Nek2 knockout U2OS cells. The data provided collectively explain the typical process by which PLK4 triggers an increase in centrosomes and define the formation of structures called CRCs, which are made up of multiple CRs connected by canonical centrosomal linkers.

## Materials and methods

### Cell lines and viral constructs

U2OS cells were cultured in DMEM medium supplemented with 10% tetracycline-free FBS (Biowest, S181T) and were periodically tested for mycoplasma contamination. To induce PLK4 expression, doxycycline hyclate was added to the medium at a concentration of 2 μg/ml. The doxycycline containing DMEM medium was refreshed every 24 hours during the experiments.

The PLK4-GFP expression construct was obtained as a gift from Michel Bornens (Addgene plasmid, 69837). FLAG-PLK4 cDNA (CDS) was cloned into the lentiviral pCW57-hygro plasmid (Addgene plasmid, 80922) by double digestion with NheI and BamHI and ligation with T4 ligase. The retroviral Centrin2-GFP expression plasmid was kindly provided by YIain Cheeseman (Addgene plasmid, 69745), while the retroviral Nek2 plasmid was acquired from DNASU (Backbone: pJP1520, Clone ID: FLH181120.01X).

The lentiviral plasmids were packaged with psPAX2 and pVSVG plasmids, and the retroviral plasmids were packaged with pUMVC and pVSVG in HEK293T cells. Transfections were performed with FuGENE according to the manufacturer’s protocol (Promega; E2311). The cell culture medium was collected, filtered through a 0.45 μm filter, and lentiviral particles were concentrated 100X by PEG8000 (Sigma; 89510). U2OS cells were transduced with the viruses in the full growth medium containing protamine sulfate (8 μg/ml). For the selection of transduced U2OS cells, puromycin (2 μg/ml), blasticidin (20 μg/ml), and hygromycin (250 μg/ml) were used. All viral-transduced cells were cultured in DMEM medium containing the selective antibiotic at a 5% concentration of the selection dose.

### CRISPR/Cas9 guided knockout

LentiCRISPR-system [18] was used to perform CRISPR/Cas9 targeted knock-out of C-Nap1, Rootletin and Nek2. Specific sgRNA’s were designed in Benchling (https://www.benchling.com). Forward and reverse oligonucleotides were obtained from Macrogen Europe BV (The Netherlands) and the sequences used were as follows: C-Nap1-sgRNA, forward: 5’-CACCGCGGCTGCAGAAGCTCACTG-3’, reverse: 5’-AAACCAGTGAGCTTCTGCAGCCGC-3’; Rootletin-sgRNA, forward: 5’-CACCGAATGGCGAGCTCATCGCGCT-3’, reverse: 5’-AAACAGCGCGATGAGCTCGCCATTC-3’; Nek2-sgRNA, forward: 5’-CACCGACATCGTTCGTTACTATGAT-3’, reverse: 5’-AAACATCATAGTAACGAACGATGTC-3’. The annealed oligonucleotides were then cloned into the LentiCRISPRv2 vector via BsmBI digestion and T4 ligation.

Lentiviruses were produced and transduced into U2OS cells. Following puromycin selection, the survivor cells were seeded at a density of 0.6 cells per well into 96-well plates, and single cell clones were expanded. The clones were monitored via Western blotting, and the single cell clones that had completely lost the expression of the target protein were used in further experiments.

### Immunofluorescence microscopy and antibodies

Cells were grown on coverslips and fixed with ice-cold methanol at -20°C for 10 min. The coverslips were then blocked with %5 BSA (Sigma; A3733) and primary antibodies were diluted in %1 BSA and incubated overnight at 4°C. The secondary antibody incubations were carried out at room-temperature for 1 hour, and washes were performed with PBS. Then coverslips were dried and then mounted on glass slides using a mounting medium containing DAPI (Vector Laboratories; H-1000-10).

The primary antibodies and their concentrations used in the experiments were: Centrin-3 (Abnova; H00001070-M01) at1/500 dilution; γ-tubulin (Sigma; T6557) at 1/500 dilution; γ-tubulin (Sigma; T3559): 1/250; CEP120 (Atlas Antibodies; HPA028823): 1/500; CEP152 (Bethyl Labs; A302-479A) at 1/500 dilution; CEP164 (Proteintech; 22227-1-AP) at 1/500 dilution; CEP170 (Bethyl Labs; A301-024A) at 1/250 dilution, C-Nap1 (Millipore; MABT1353) at 1/500 dilution; CEP68 (Proteintech; 15147-1-AP) at 1/500 dilution; Rootletin (Santa Cruz Biotechnology; sc-374056) at 1/500 dilution; Nek2 (BD; 610593) at 1/500 dilution. The secondary antibodies used in the experiments were: anti-rabbit AF488 (Invitrogen; A-11008), anti-rabbit AF555 (Invitrogen; A-31572), anti-mouse AF594 (Abcam; ab150116) and anti-mouse AF647 (Abcam; ab150115).

### High resolution confocal microscopy

The Leica DMi8 microscope was used for confocal microscopy. All images were captured using a 100x objective (HC PL Apo CS2 100x / 1.40 OIL). Huygen’s deconvolution was performed using the following parameters for each channel: Minimum iterations: 40; Signal to noise ratio: 20; Quality threshold: 0.05; Iteration mode: optimized; Brick layout: auto; Vertical mapping function: logarithmic; Estimation mode: automatic; and Area(radius): 0.7 μm. The deconvolved images were imported to LASX (Leica) software, and 3D reconstructions were generated from the deconvolved images. All Z-stack images used in the study were acquired as 0.2 μm slices, and maximum projections of the Z-stacks were used to generate all 2D images.

For counting and distance measurement experiments, a 63x objective was used, and images were analyzed with LASX. To measure the distances between CRs, the centers of the two γ-tubulin foci were connected with a line, and the length of the line was measured. To measure the diameter of the CRC, a circle was drawn around the CRC structure, and the diameter of the circle was measured as seen in Fig. S3C.

### Cell cycle synchronization and flow cytometry

Aphidicolin and double thymidine block (DTB) were used in cell cycle synchronization experiments. 1.6 μg/ml aphidicolin (Sigma; A0781) was added to the medium for 16 hours and cells were washed and released into full growth medium. When S-Trityl-L-cysteine (STLC) was used after aphidicolin block, cells were released in normal medium for 2 hours, then incubated with 5 μM STLC (Cayman Chemical; 23236) for 8 hours. DTB was performed with the addition of 2.5 mM thymidine (Sigma; T9250) to full growth medium for 18 hours, and block was performed 2 times. Cells were released into full growth medium for 8 hours between the 2 thymidine blocks. Cell pellets were fixed by ethanol in -20 °C, washed with PBS, treated with 100 μg/ml RNAse A (Thermo; 12091021), and stained with 50 μg/ml propidium iodide solution (Sigma; P4170). The cell cycle distribution of cell populations was analyzed with a flow cytometer (Cytoflex, Beckman Coulter). Generated .fcs files were analyzed and visualized by FlowJo (v10.8.1).

### qPCR and Western blotting

RNA was extracted from cells using Nucleospin RNA (Macherey-Nagel; 740955) and cDNA was synthesized from 1 μg of total RNA using the M-MLV (Invitrogen; 28025013). SYBR green master mix (Roche; 04707516001) was used to amplify 10 ng of cDNA template. The primers used were as follows: PLK4, forward: 5’-GGCCAAGGACCTTATTCACCA-3’, reverse: 5’-TGTGGCATGCCCACTATCAA-3’; β-actin, forward: 5’-AGCACAGAGCCTCGCCTT-3’, reverse: 5’-CATCATCCATGGTGAGCTGG-3’. Real-time quantitative PCR was performed on a LightCycler 480 (Roche), and the relative fold change in gene expression was measured with the 2^-ΔΔCT^ method.

SDS-PAGE and Western blotting were performed following standard protocols. The following primary antibodies were used: Anti-FLAG (Sigma; F3165), Anti-C-Nap1 (Santa Cruz Biotechnology; 390540), Anti-Rootletin (Santa Cruz Biotechnology; sc-374056), Anti-Nek2 (BD; 610593), Anti-GAPDH (Abcam; ab9485), and Anti-β-actin (Abcam; ab8227). Immunoreactive bands were developed with Luminata Forte (EMD Millipore) and visualized in an Odyssey FC imaging system (Licor).

### Statistical analyses

Data from multiple groups were compared by One-way ANOVA Dunnett’s Multiple comparisons test. p value smaller than 0.05 was considered to be statistically significant. Frequency distributions were calculated with non-linear gaussian regression.

## Results

### Long term PLK4 induction generates centrosome rosette clusters

The structure and protein composition for centrosome rosettes (CRs) have been characterized in several previous research articles which have contributed to a better understanding of centriole biogenesis processes [19] and chromosomal defects in cancer [16]. However, to our knowledge, the cell cycle dependent maturation of CR structures and how long-term PLK4 induction affects centrosome structures have remained unexplained. Here, we first set out to observe the appearance of CRs in high resolution. We over-expressed a GFP-tagged PLK4 construct in U2OS cells for 24 hours and used confocal microscopy followed by deconvolution for imaging of the CR structures. As expected, short term over-expression of PLK4 resulted with cells that contained two CRs (Fig. 1A) 1.

**Fig 1.**
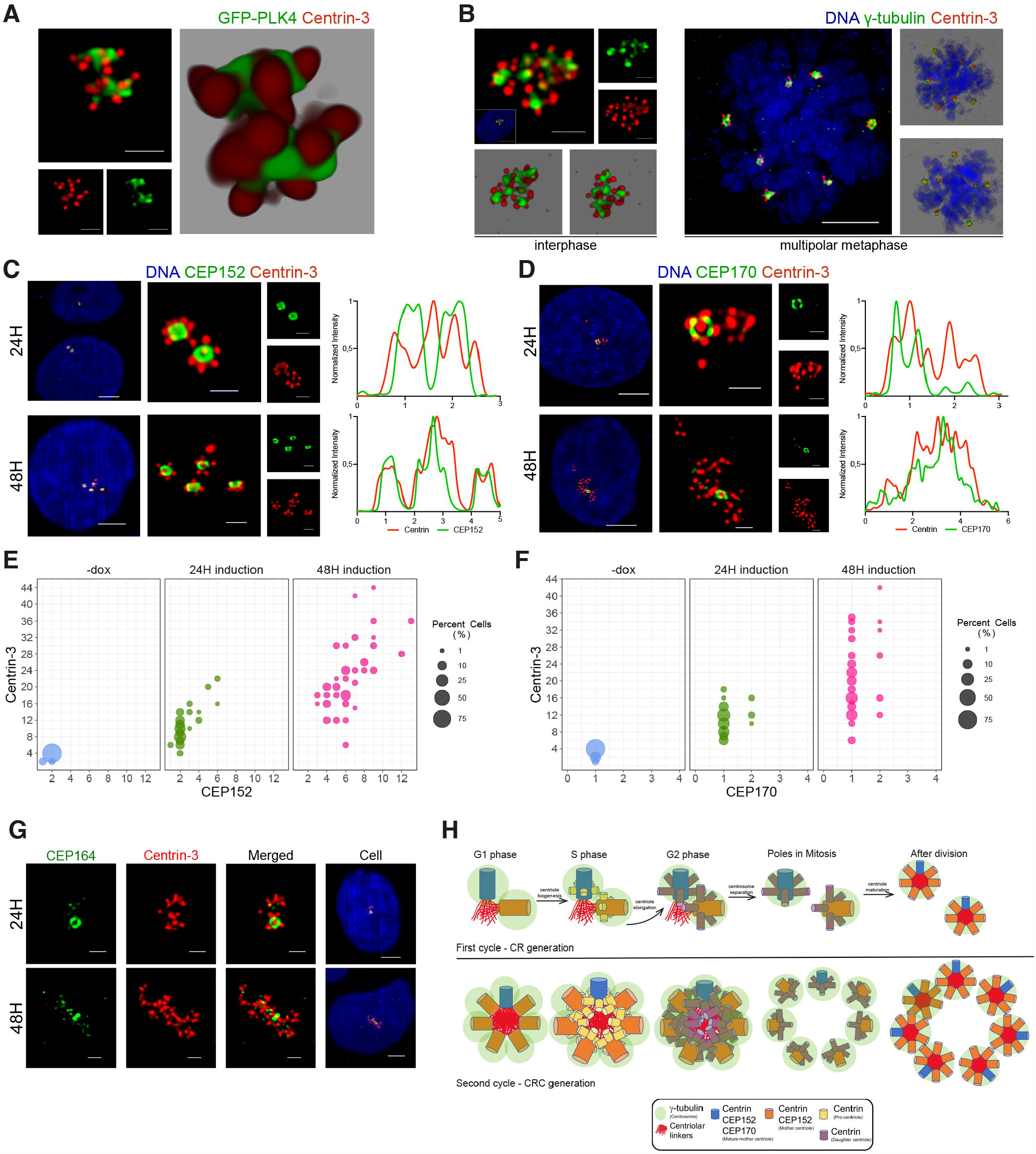
Long term induction of PLK4 leads to the formation of CRCs. A) Over-expression of PLK4 for 24 h generates cells with 2 centrosome rosettes. U2OS cells were transfected with GFP-tagged PLK4 (green) and stained for Centrin-3 (red). The left panel shows a high-resolution confocal image, and the right panel shows a 3D reconstruction of the confocal image. B) Induction of PLK4 for 48 h generates cells with multiple CRs. PLK4 expression was induced with dox (48 h), and cells were stained with DAPI (DNA, blue), γ-tubulin (centrosome, green) and Centrin-3 (centriole, red). The left panel shows an interphase cell with a CRC, and the right panel shows multipolar metaphase formation with CRs in each pole. C) The localization of CEP152 (green) and Centrin-3 (red) in PLK4-induced cells for 24 h (upper panel) and 48 hours (bottom panel). D) The localization of CEP170 (green) and Centrin-3 (red) in PLK4-induced cells for 24 h (upper panel) and 48 hours (bottom panel). The right panels show normalized fluorescence intensities in C and D. E) The quantification of mother centrioles and total centriole numbers in PLK4-induced cells for 24 h and 48 h. n:100, N:2 biological repeats. F) The quantification of mature-mother centrioles and total centriole numbers in PLK4-induced cells for 24 h and 48 h. n:100, N:2 biological repeats. Raw counting data of 1E and 1F is available in Supplemental Table 1. G) The localization of CEP164 (green) and Centrin-3 (red) in PLK4-induced cells for 24 h (upper panel) and 48 h (bottom panel). H) A model for the generation of CRCs. A newly divided G1 cell has two centrosomes containing one mature-mother and one young mother centriole. Increased PLK4 levels generate >1 procentriole on the wall of mother centrioles at the beginning of S phase, leading to the generation of CRs. When a cell with CRs divides into two daughter cells, each daughter cell receives one CR. Daughter centrioles in a CR get disengaged after division, mature to become young mother centrioles, and are linked to other mother centrioles with centriolar linker proteins. When PLK4 induction continues, all mother centrioles generate linked CRCs.

In line with predictions, following the induction of PLK4 for 24 h in U2OS cells that contain a doxycycline (dox) inducible promoter (Fig. S1A), CR structures were readily detected in interphase cells (Fig. S1B). In order to understand how CRs would appear if they over-duplicate more than once (2 cell cycles), we performed long-term PLK4 induction (48 hours), and this resulted in centrosome amplification, similar to previously reports [13, 16, 20, 21], as evidenced by the presence of >2 γ-tubulin foci per cell (Fig. S1C). Interestingly, amplified centrosomes were clustered in most of the interphase cells and each had a CR (Fig. 1B and S1C). Since they were located in close proximity, we hypothesized that CRs may be connected with centrosomal linker proteins and named these structures as “centrosome rosette clusters” (CRCs). Furthermore, we observed the formation of multipolar metaphases with a single CR at the spindle poles, indicating that individual CRs acted similar to single centrosomes (Fig. 1B) 1. When the number of γ-tubulin and Centrin-3 foci were scored in cells following 24- and 48-hours induction of PLK4, centriole numbers were increased first (24 h) with no change in centrosome number, which was elevated dramatically after 48 h (Fig. S1D).

We utilized various markers to determine the level of maturation of amplified centrioles. Centrin-3 was used to label all centrioles at different stages of development, from pro-centriole to mature-mother centriole [22]. CEP152 was used to identify the proximal region of the mother centriolar wall, appearing as a ring around the mother centrioles [23, 24]. Distal and subdistal appendage proteins, CEP164 and CEP170, respectively, were used to mark the fully matured mother centriole during interphase [25]. CEP120 was employed to identify the centriole wall, primarily of daughter centrioles [26].

In order to ensure the specificity of our antibodies, we first performed co-staining of U2OS cells expressing GFP-Centrin-2 with γ-tubulin and CEP120. Our findings demonstrated that CEP120 staining had 4 foci per cell in cells with duplicated centrosomes, and the staining was localized in newly generated daughter centrioles. Additionally, we detected the presence of two ring-shaped structures through CEP152 staining, which co-stained with 2 γ-tubulin foci. Conversely, CEP164 and CEP170 antibodies only labeled one mother centriole, the mature-mother centriole, as shown in Fig S1E. These morphological observations are consistent with the anticipated distribution patterns of the proteins.

To explore the effects of PLK4 over-expression on centriole amplification and maturation, we performed staining using Centrin-3 and CEP152 (Fig 1C, Fig S1F) 1 or CEP170 (Fig 1D, Fig S1G) 1 after 24 and 48 hours of PLK4 induction. The findings revealed that 24 hours of PLK4 induction resulted in the presence of numerous cells with two centrosome rosettes (CRs). However, following 48 hours of induction, the number of CEP152 rings per cell increased, indicating the amplification of centrosomes. Additionally, the number of Centrin-3 foci per cell has also increased after 48 hours of induction, suggesting that high levels of PLK4 caused another round of centriole amplification on previously amplified mother centrioles (Fig 1E) 1. CEP170 staining was positive in only one CR after 24 hours and one or two CRs in 48 hours PLK4 induction, suggesting mature-mother centriole content is not changed as CEP152 foci with the duration of PLK4 induction (Fig 1F, Fig S1G) 1. In 24 hours of PLK4 induced cells, CEP120 only co-stained with Centrin-3 signals from surrounding centrioles of a CR, providing evidence that the surrounding Centrin-3 signals were newly generated daughter centrioles and leaving the centriole located in the middle (mother centriole) unstained (Fig S1H). Additionally, CEP164 staining also showed a similar pattern to CEP170 in both 24 and 48 hours of PLK4 induced cells (Fig 1G) 1. These observations are consistent with previous studies in the field [16], indicating that while prolonged PLK4 induction causes centrosome amplification, the mature-mother centriole content of a cell is not altered to an excessive number of CEP152-positive mother centrioles.

Here we propose a model of CRC formation by prolonged PLK4 induction based on our observations (Fig 1H) 1. A cell in G1 phase has two centrosomes, consists of one centriole each, a mature-mother and a mother. At the end of the G1/early S phase, a cell has two centrosomes, each consisting of a mature-mother and a mother centriole. In the first cell cycle following PLK4 induction, CRs are generated that function as single centrosomes. During cell cycle progression, the newly generated daughter centrioles elongate and dissociate from their mother centrioles during mitosis. In the following G1 phase, daughter centrioles mature into mother centrioles capable of recruiting PLK4 and PCM proteins such as CEP152 and γ-tubulin. Upon entry into S phase, high levels of PLK4 lead to centriole amplification on each mother centriole, resulting in CRC formation. The model also explains the low number of mature-mother centrioles in cells with CRCs because a daughter cell inherits only one mature-mother centriole from its parent. It is noteworthy that this model of CRC formation requires linking all CRs with centrosomal linkers until separation in G2/M phase.

### CRs and CRCs are linked with centrosomal linker proteins

Previously, to our knowledge, maturation of amplified centrioles via PLK4 induction was depicted through illustrations which suggested that separate CRs were generated without any functional linkage between amplified centrosomes [16, 27]. In our proposed model (Fig 1H), we suggest that PLK4 induced CRCs should be linked to each other with centrosomal linker proteins until separation in prophase. In order to obtain evidence to support this, we first investigated the localization of Rootletin, CEP68 and C-Nap1 in cells with 24 and 48 hours PLK4 induction.

Rootletin is a self-assembling filamentous protein that connects interphase centrosomes by creating spider web like filaments [28, 29]. CEP68 is another centrosomal linker protein that binds and organizes Rootletin fibers into thick filaments [30]. C-Nap1, on the other hand, is a proximally localizing centriolar protein that assembles as a ring and functions as an anchor for centrosomal linker proteins [30, 31].

As anticipated, our findings demonstrated that Rootletin and CEP68 bind interphase CRs in cells induced with PLK4 for 24 hours (Fig. 2A and 2B) 2. After 48 hours of dox induction, the CRCs were also interconnected with centrosomal linkers. Strikingly, we noticed that a significant number of cells exhibited a circular arrangement of CRCs that were linked together by a ring-shaped centrosomal linker network composed of Rootletin (Fig. 2C, Fig. S2A) 2 and CEP68 (Fig. 2D, Fig S2B) 2. To confirm the localization of Rootletin between the CRs and CRCs in higher resolution, we used STED microscopy (Fig. S2C). Co-staining of Rootletin and CEP68 revealed similar localization patterns of both proteins in CRCs (Fig S2D). We also found that all mother centrioles in cells with CRs and CRCs were positive for C-Nap1 (Fig. 2E) 2, as well as the primary centrosome separator protein Nek2(Fig. 2F) 2.

**Fig 2.**
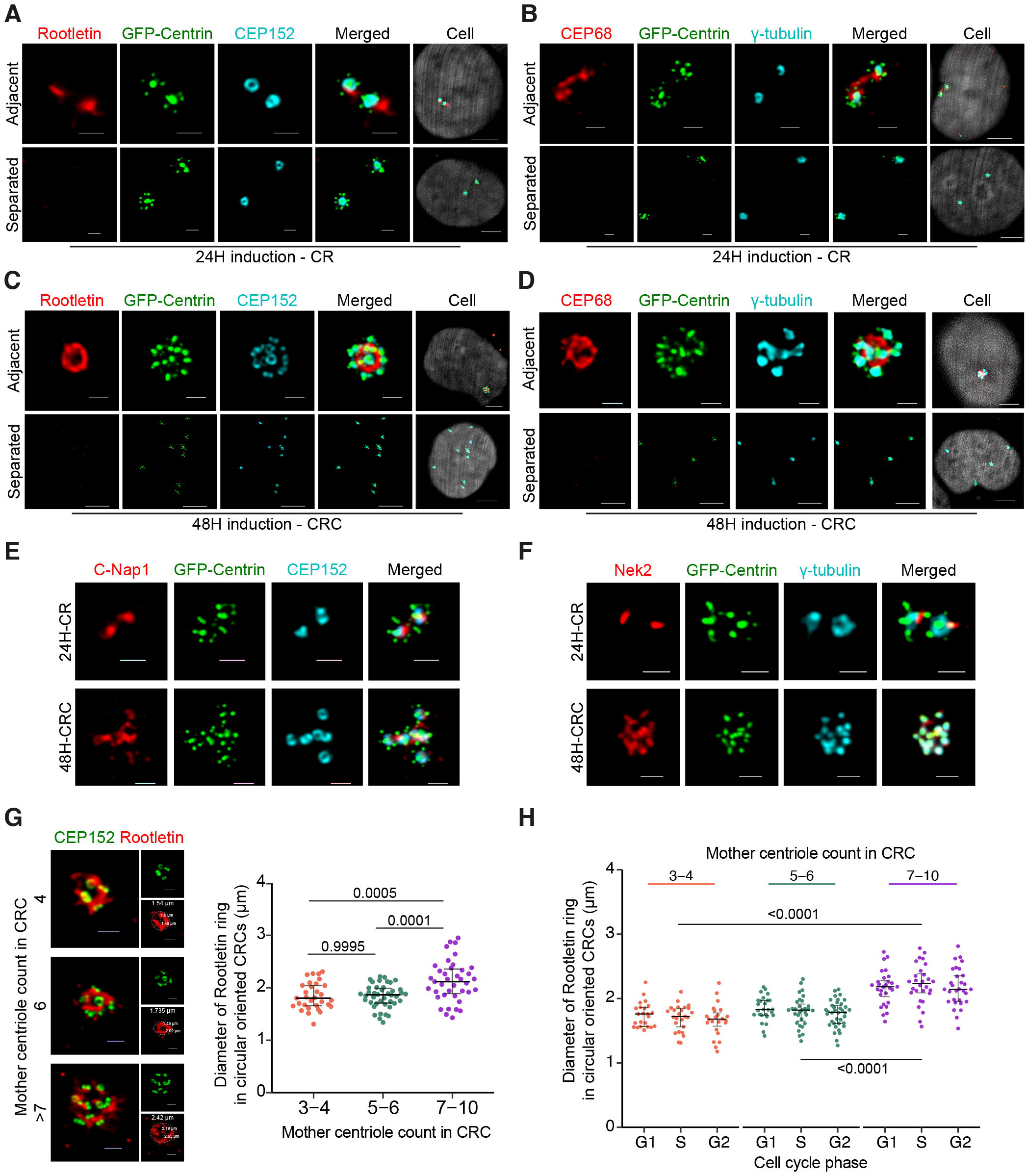
Centrosomal linker proteins connect CRs and CRCs. A-B) Rootletin and CEP68 binds adjacent CRs in 24 hours PLK4 induced cells. Scale bars: single staining and merged channels: 1 μm, Cell: 5 μm. C-D) Rootletin and CEP68 binds CEP152 positive mature centrioles of CRCs in 48 hours PLK4 induced cells. Scale bars in adjacent groups: single staining and merged channels: 1 μm, Cell: 5 μm. Scale bars in separated groups: 5 μm. E-F) Mother centrioles in CRs and CRCs are positive for C-Nap1 (E) and Nek2 (F). Scale bars: single staining and merged channels: 1 μm, Cell: 5 μm. G) Left panel: Representative images of circular linked CRCs with different number of mother centrioles. Scale bars: 1 μm. Right panel: Diameter of Rootletin ring in circular oriented CRCs are increased in cells with >7 mother centrioles (n: 112 circular linked CRC from 2 independent experiment). H) Diameter of Rootletin ring in circular linked CRCs is not changed with cell cycle progression (n; G1: 77, S: 84, G2: 90 circular linked CRCs from 2 independent experiments.). Lines represent median and interquartile range in G and H. Raw measurement data of 2G and 2H is available in Supplemental Table 2

We observed two types of centrosomal linker orientation in cells with CRCs; (i) circular oriented, which is characterized with arrangement of mother centrioles around a circular shaped linker, and (ii) planar oriented, that mother centrioles are distributed around and bound with linear linker in between (Fig. S2E). When scored, 45% of the CRCs exhibited a circular arrangement, while the rest were arranged in a planar fashion (Fig. S2F). To gain a better understanding of the circular linker arrangement, we measured the diameter of the circular linkers. The mean diameter of the circular-oriented linkers was 1.92 μm (Fig. S2G). Interestingly, the diameter of the Rootletin ring did not change based on the number of mother centrioles surrounding the linker in CRCs with 3-6 mother centrioles, but it significantly increased in CRCs with more than 7 mother centrioles (Fig. 2G) 2.

We also investigated whether the binding and spatial positioning of amplified centrosomes were regulated by the cell cycle. However, quantification of CRC linkage phenotypes in synchronized cells showed that the percentage of observed linkages did not change with cell cycle progression, indicating that the observation of the two different types of linkage orientation in CRCs was a distinct phenotype, not a cell cycle-dependent event (Fig. S2H). We also found that the diameter of the circular Rootletin linker remained constant across all cell cycle phases and was not affected by cell cycle progression. As in unsycnhronized cells (Fig. 2G) 2, the diameter of the circular linker was significantly higher in CRCs with more than 7 mother centrioles than CRCs with 3-6 mother centrioles in all cell cycle phases (Fig. 2H) 2.

In addition, separation of CRs and CRCs resulted with reduced Rootletin and CEP68 staining intensity (Fig. 2A-D) 2, suggesting linker components of CRCs are functional and regulated as normal centrosomes. Furthermore, other centrosomal linker components, such as LRRC45, were found to be present in the inter-CRC linker [32] (Fig. S2I). Collectively, these findings provide evidence for our CRC generation model and indicate that amplified centrosomes induced by PLK4 are connected to each other through canonical centrosomal linkers.

### C-Nap1 (CEP250) and Rootletin (CROCC) knockout results in distanced CRCs

As CRs and CRCs are connected with centrosomal linker proteins, and the staining intensity of the linker was decreased in separated CRs and CRCs, we hypothesized that the regulation of CRs and CRCs is influenced by mechanisms related to centrosome linking and separation. To investigate this, we focused on three specific proteins: C-Nap1, which acts as an anchor for centrosomal linker proteins; Rootletin, the primary centrosomal linker; and Nek2, an important regulator of centrosome splitting [33]. We discovered that knocking out any of these proteins individually using the CRISPR/Cas9 system (Fig. S3A) did not have any impact on the generation of CRs and CRCs induced by PLK4 expression (Fig. 3A) 3. Additionally, after 72 hours, the levels of centrosome amplification were comparable in all KO cell clones, indicating that the absence of these proteins did not hinder PLK4-induced centrosome amplification (Fig. S3B). Therefore, it appears that these proteins are not required for the formation of CR and/or CRC.

**Fig 3.**
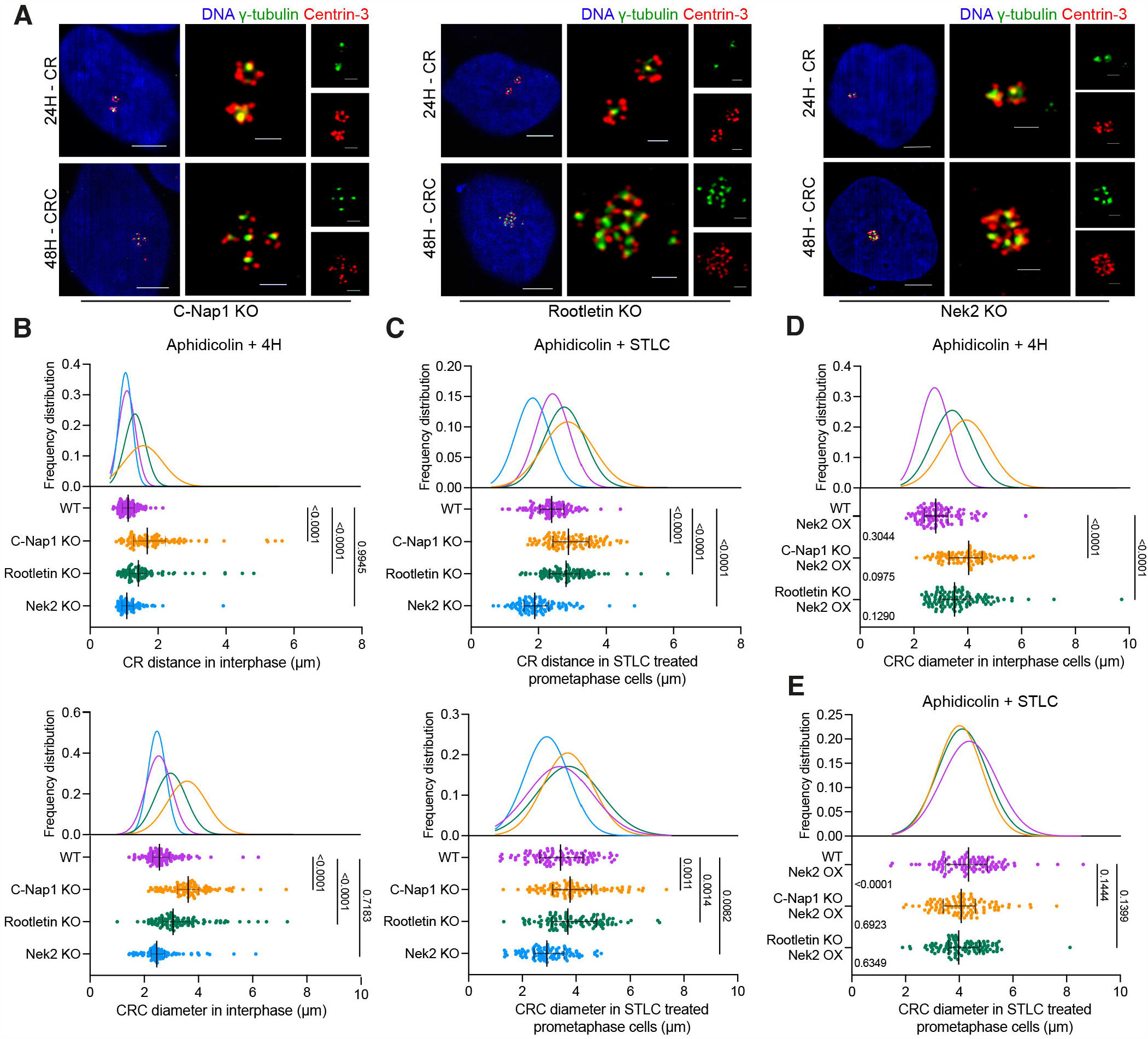
The regulation of CRC binding and separation involves centrosomal linkers and Nek2. A) CR (top panel) and CRC (bottom panel) formations in C-Nap1 KO (left panel), Rootletin KO (middle panel) and Nek2 KO (right panel) U2OS cells. B) CR distances (top panel) and CRC diameters (bottom panel) in cells synchronized with aphidicolin and released for 4 hours. n: 100 for each group, pooled from 4 independent experiment. C) CR distances (top panel) and CRC diameters (bottom panel) in prometaphase cells synchronized with aphidicolin and STLC. n: 100 for each group, combined from 4 independent experiment. D) CRC diameters of Nek2 over-expressing WT, C-Nap1 KO and Rootletin KO cells in interphase. Left side p values indicate comparisons with control groups in Fig. 3B bottom panel. n: 100 for each group, pooled from 3 independent experiment. E) CRC diameters in Nek2 over-expressing WT, C-Nap1 KO and Rootletin KO cells in prometaphase. Left side p values indicate comparisons with control groups in Fig. 3C bottom panel. n: 100 for each group, combined from 3 independent experiment. B-E: Median and interquartile range is shown on plots. Frequency distributions are calculated with non-linear gaussian regression and raw measurement data is available in Supplemental Table 3.

Then, to further explore the role of centrosomal linkers in CRCs, we examined the distancing of CRCs in our KO cell models. We found that the diameter of CRCs in interphase cells were increased in C-Nap1 and Rootletin knock-out cells, while Nek2 knock-out had no impact, suggesting that the linkers play an essential role in both the formation and positioning of CRCs (Fig. S3C). Since the centrosome cycle is regulated by the cell cycle, we also evaluated the centrosome distancing phenotype in synchronized populations. Aphidicolin and double thymidine block (DTB) were used to block cells in S and G1 phases. After synchronization, aphidicolin-synchronized cells were released into full growth medium for 4 hours to allow cells to progress through S phase (Fig. S3D). Similar to unsynchronized cells, the distances between CRs and the diameters of CRCs in C-Nap1 and Rootletin KO cells were increased (Fig. 3B) 3. Likewise, G1-arrested cells were released for 6 hours, resulting in S-G2-enriched cell populations (Fig. S3D), and the results were comparable to those of aphidicolin-synchronized samples (Fig. S3E). In both experimental approaches, Nek2 KO did not affect the distancing of interphase CRs and CRCs. In summary, the loss of centrosomal linkers led to the distancing of interphase CRCs.

### Nek2 regulates pre-mitotic separation of CRCs

Centrosome separation is primarily regulated by Nek2 serine/threonine kinase and Eg5 mitotic kinesin [34]. During prophase, Nek2 phosphorylates various centrosomal linkers, including C-Nap1, Rootletin, CEP68, and LRRC45, to regulate centrosome separation [32, 35–37]. Meanwhile, Eg5 kinesin is responsible for the main mitotic microtubule motor force that separates centrosomes [38, 39]. Since canonical centrosomal linkers connect the CRCs (Fig. 2H) 2, we hypothesized that Nek2 should regulate the pre-mitotic separation of PLK4-induced CRCs. To investigate this, we inhibited Eg5 kinesin using S-trityl-L-cysteine (STLC) in our KO cell clones and evaluated CR and CRC distances in prometaphase cells (Fig. S3D, S3F and S3G). We found that CRs and CRCs that were already distanced in interphase (Fig. 3B) remained distanced in prometaphase in C-Nap1 and Rootletin KO cells (Fig. 3C). In contrast, both CRs and CRCs in Nek2 KO cells remained adjacent, indicating that Nek2 is necessary for pre-mitotic separation of CRCs in cells with inhibited Eg5 kinesin (Fig. 3C) 3.

Additionally, we over-expressed Nek2 in U2OS cells that were either wild type, C-Nap1 KO, or Rootletin KO (Fig. S3H) in order to assess: (i) how increased levels of Nek2 affect the separation of CRCs, and (ii) whether centrosomal linkers are necessary for Nek2-mediated CRC separation. Our findings showed that elevated levels of Nek2 did not significantly impact the separation of CRCs in interphase cells that were synchronized with aphidicolin (Fig. 3D) 3. We then utilized STLC to inhibit Eg5 kinesin and assessed the distances between CRCs in prometaphase cells. The results demonstrated that Nek2 over-expression led to an increase in CRC distancing only in wild-type cells, and not in cells lacking C-Nap1 or Rootletin (Fig. 3E) 3. Therefore, our data suggests that Nek2 plays a role in regulating the separation of amplified centrosomes induced by PLK4 via acting on centrosomal linkers.

### PLK4 induction generates CRCs during second cell cycle

To better understand how CRs and CRCs progress through the cell cycle, we synchronized U2OS^dox-PLK4^ cells using DTB and monitored them for 32 hours at 4-hour intervals (Fig 4A) 4. We selected six time points (0h, 4h, 8h, 16h, 24h, 28h) that best reflect the consecutive phases of two cell cycles (G1-1, S-1, G2-1, G1-2, S-2, G2-2) after PLK4 induction. We then stained the cells for CEP152 and Centrin-3 to visualize CRs and CRCs and observed that CRs were generated in the 4h and 8h samples, and most cells formed CRCs in the 16h, 24h, and 28h samples (Fig. 4B) 4. The centriole number per cell increased in the first cell cycle (4h, 8h) and the number of CEP152-positive mother centrioles per cell increased in G1-2 (16h). After cell division and progression through S-2 and G2-2 phases, most cells contained more than 10 centrioles and more than 4 mother centrioles (Fig. 4C) 4.

**Fig 4.**
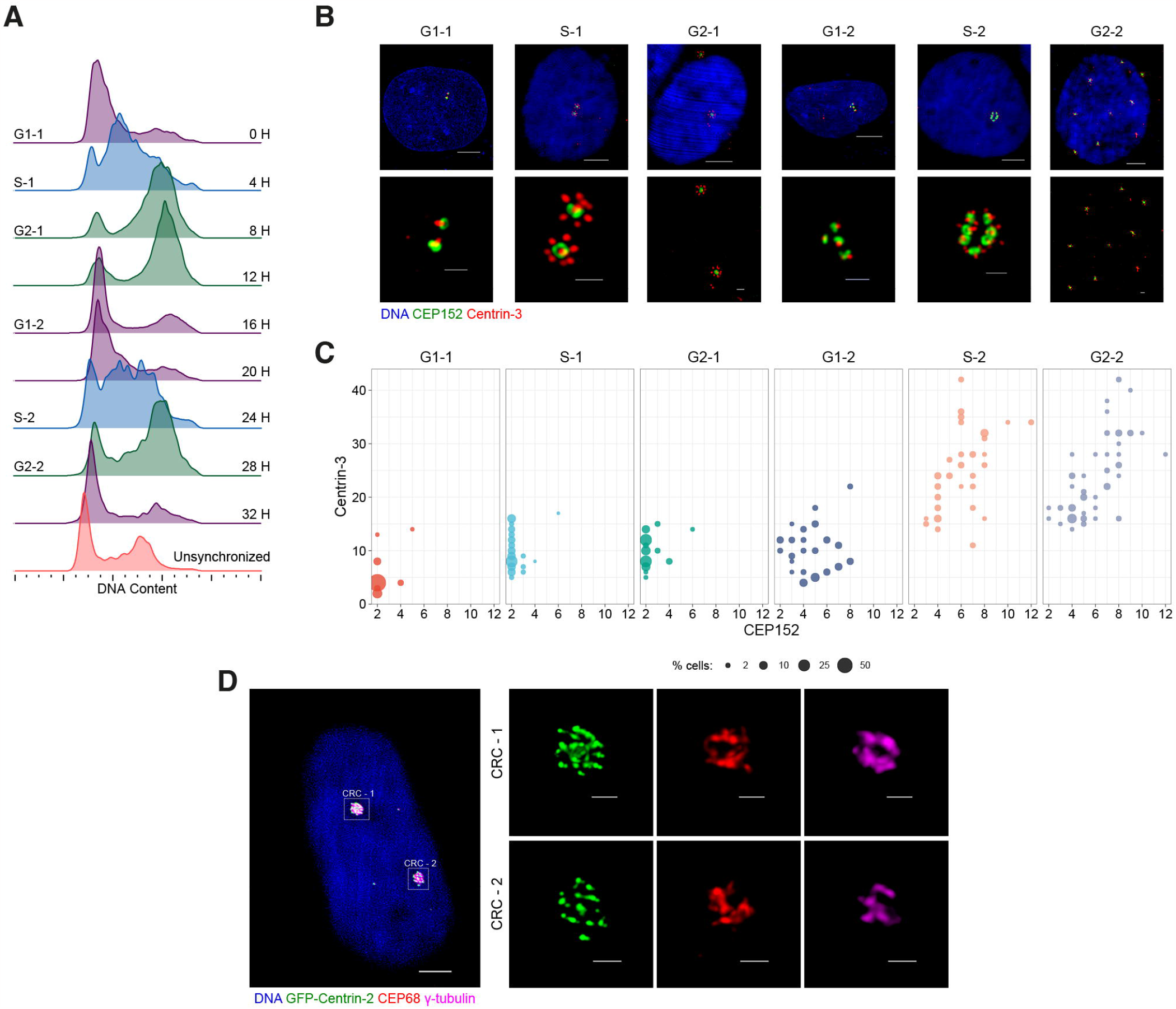
Centrosome rosette clusters develop during the second cell cycle following PLK4 induction. A) Cell cycle synchronization using double thymidine block (DTB). B) Representative images of CRs and CRCs in cells at various cell cycle phases. C) Quantification of mother centrioles and total centrioles in cells with CRs and CRCs across different cell cycle stages. D) A representative image of a cell containing two distinct CRCs in 36h PLK4 induced cells.

Next, we focused on cells in the third cell cycle (36h after DTB release) after PLK4 induction, which are the progeny of cells that divided with centrosome clustering in the previous cycle. To identify cells that generated more than one CRC, we stained GFP-Centrin2 expressing U2OS cells for CEP152 and Rootletin and observed the presence of several cells with two independently inter-connected CRCs (Fig. 4D) 4.

In summary, our results show that CRs are formed in the first cell cycle after PLK4 induction, mature and generate CRCs in the second cell cycle, and cells can generate more than one CRC in subsequent cell cycles, providing support for our model as illustrated in Fig. 1H.

## Discussion

Centrosome amplification is now widely recognized as a significant characteristic of cancer [40], and increased expression of PLK4 in human tumors is responsible from centrosome amplification in cancer [20, 41–43]. Our research contributes to this knowledge by demonstrating the binding of PLK4-induced amplified centrosomes through centrosomal linkers, providing a mechanistical insight to PLK4-induced centrosome amplification.

Recent studies have revealed that PLK4 induced centrosome amplification is not only present in cancer cells but is also necessary for generating multiciliated cells [17, 44]. Previous research demonstrated that inducing PLK4 expression in human cells leads to centrosome rosettes (CRs) in the first cell cycle [19], followed by centrosome amplification in the second cell cycle [8]. In line with these findings, our study shows that PLK4 induction for 24 hours generates cells with CRs, while 48 hours of induction results in CRCs or centrosome amplification. Notably, after 48 hours of induction, we observed an increase in the number of CEP152 positive mother centrioles in the cell, but the number of CEP164 and CEP170 positive mature mother centrioles remained comparable to cells with normal centrosomes. This perfectly fits with our model in Fig.1H, which suggests PLK4 induction would generate only one CR which contains a mature mother centriole. Additionally, U2OS cells employed in our research have wild-type p53, which activates the PIDDosome pathway and triggers apoptosis in cells with a high number of mature mother centrioles [45]. This pathway could account for the low percentage of cells with more than two mature mother centrioles observed after 48 hours of PLK4 induction. In future studies, it would be intriguing to investigate the development and maturation of CRCs in a population of cells with a mutated p53 gene.

Our study provides important insights by demonstrating that the amplified centrosomes in cells during the second cell cycle after PLK4 induction are indeed connected via centrosomal linkers, as we hypothesized in Fig 1H 1. Additionally, we confirmed the functionality of these linkers in CRCs and provided evidence that the separation of PLK4-induced amplified centrosomes requires Nek2 activity, similar to the separation of a normal pair of centrosomes. It is noteworthy that we have noticed that CRCs can adopt either a planar or circular binding, with their occurrence almost equally split between the two shapes. When the orientation is planar, the CRCs do not have a particular form. However, when the orientation is round, there appears to be a constraint, with a maximum of six centrioles encircling the mother centriole, where bridging via centrosomal linkers can take place without an increase in diameter. This suggests that the maximum capacity for a well-organized CRC structure may be limited to six mother centrioles, beyond which the diameter of the linker increases, possibly due to the inability to accommodate or connect additional mother centrioles. Based on the average diameter of a centriole, which is approximately 0.2-0.5 μm [46, 47], we can estimate that a circular arrangement of six centrioles without any gaps between them would have a circumference of about 1.2-3.0 μm and a diameter of about 0.4-1 μm. However, this is smaller than the observed average diameter of 1.92 μm, which suggests that the additional mother centrioles are not arranged adjacent to one another, but rather with at least one centriole distance in between them. A circular structure with more than six centrioles is still possible, albeit with an increase in the diameter.

Therefore, these distinct structures enable specific alterations in shape while maintaining the morphology of the CRCs. Our results revealed that absence of centrosomal linker proteins resulted with increased distances between CRs and CRCs, however, CRCs were still observed in close vicinity. Given that actin forces and kinesin motor proteins as KIFC3 are also important in centrosome positioning [48], knocking-out centrosomal linker proteins individually may show a limited effect.

The over-expression of PLK4 is a double-edged sword for cancer cells, as it can provide an advantage in promoting genomic instability, while also having the potential to form multipolar metaphases due to extra centrosomes, which can result in uneven cell division. Since such divisions may eventually cause the loss of essential genetic content, cancer cells typically evade the negative effects of multipolar divisions by generating pseudo-bipolar spindles [27]. Numerous studies have shown that the formation of pseudo-bipolar spindles is a major cause of chromosome missegregation and chromosomal instability [8, 49]. Given the extensive range of centrosome clustering mechanisms discovered [50], it is crucial to further explore how cancer cells can effectively cluster PLK4-induced amplified centrosomes to prevent multipolar cell divisions.

In this study, we presented evidence that PLK4-induced extra centrosomes cluster and are linked through canonical centrosomal linker proteins, similar to normal centrosome pairs. Furthermore, we demonstrated that CRCs are regulated by cell cycle progression and canonical centrosome linking and separation mechanisms. Considering that centrosome amplification and mutations in linker protein-coding genes are associated with numerous diseases, we believe that comprehending the mechanisms behind the binding and separation events of amplified centrosomes will enhance our understanding of the biology underlying PLK4-induced centrosome amplification.

## Conclusion

This study provides valuable insights into the mechanisms behind PLK4-induced centrosome amplification in cancer. By demonstrating the linking of amplified centrosomes induced by PLK4 via centrosomal linkers and the functionality of these linkers in CRCs, the study enhances our understanding of the biology underlying PLK4-induced centrosome amplification. Given that centrosome amplification and mutations of linker protein coding genes are associated with many diseases, we believe that understanding the mechanism of binding and separation events of amplified centrosomes will contribute to better understanding of the biology of PLK4-induced centrosome amplification.

## Supporting information

**S1 Fig. Supplemental to Fig. 1** A) Doxycycline induced PLK4 over-expression. B) 24-hours PLK4 induction generates CRs. Left panel: Two cells with CRs. Cells were stained with DAPI (DNA, blue), γ-tubulin (centrosome, green) and Centrin-3 (centriole, red). Right panel: Percentage of cells with >4 Centrin-3 foci in -dox and +dox conditions. C) 48-hours PLK4 induction forms CRCs. Left panel: Two cells with CRCs. Cells were stained with DAPI (DNA, blue), γ-tubulin (centrosome, green) and Centrin-3 (centriole, red). Right panel: Percentage of cells with >2 γ-tubulin foci in -dox and +dox conditions. D) Quantification of centrosomes and total centriole numbers in 24-hours and 48-hours PLK4 induced cells. n:100, N:2. E) Co-staining of CEP120 / CEP152 / CEP164 and CEP170 with γ-tubulin in GFP-Centrin-2 expressing U2OS cells. F-H) Representative images of localization of CEP152 (F), CEP170 (G) and CEP120 (H) in 24-hours (upper panels) and 48-hours (bottom panels) dox induced cells.

**S2 Fig. Supplemental to Fig. 2** A-B) 3D visualization of Z-stacks from Fig. 2C. (A) and Fig. 2D (B). C) Representative confocal and STED images showing the connection of CRs and CRCs with Rootletin. D) Representative confocal images of co-staining of Rootletin and CEP68 in cells with CRCs. E) Representative confocal images of planar oriented and circular oriented Rootletin linker in cells with CRCs. F) Percentage of CRC arrangement type in cells with CRCs. Rootletin and CEP68 staining were independently quantified; dots represent biological repeats, and lines display the mean of repeats. G) Diameter of Rootletin ring in cells that CRCs are bound with circular oriented linker. Dots represent measurement of diameter in individual cells. (n: 102 circular CRC from 2 independent experiment, line represent mean value.) H) Left panel: Cell cycle synchronization with double thymidine block, Right panel: Diameter of Rootletin ring in cells with CRCs that are bound with circular oriented linker in different cell cycle phases. G1 phase: 0h after DTB, S phase: 4h after DTB release, G2 phase: 8h after DTB release. I) LRRC45 localization in CRCs.

**S3 Fig. Supplemental to Fig. 3** A) Western blot showing PLK4 induction in C-Nap1, Rootletin and Nek2 individual knock-out U2OS cell clones. B) Centrosome amplification in 72 hours PLK4 induced U2OS-WT and U2OS-KO cells. C) Representative measurements of two cells with CRCs (blue: DAPI, green: γ-tubulin, red: Centrin-3. Note that γ-tubulin negative disengaged centrioles in bottom image are not included in diameter measurements.)(right panel) and CRC diameter in 48 hours PLK4 induced unsynchronized cells (left panel). Median and interquartile range is shown on plot. n: 50 for each group, pooled from 2 independent experiment. D) Cell cycle profiles of unsynchronized and synchronized cells. E) CR distances (top panel) and CRC diameters (bottom panel) in cells synchronized with DTB and released for 6 hours. Median and interquartile range is shown on plots. n: 100 for each group, pooled from 4 independent experiment. F) Representative images of prometaphase distances in cells with CRCs. Cells were stained with DAPI (DNA, blue), γ-tubulin (centrosome, red). G) Representative images of prometaphase distances in cells with CRCs. Cells were stained with DAPI (DNA, blue), γ-tubulin (centrosome, green) and Centrin-3 (centriole, red). H) Western blot showing Nek2 over-expression in U2OS-WT and U2OS-KO cell groups.

**S4 Fig. Uncropped Western blots** Uncropped western blot images of Fig. S1A (A), Fig. S3A (B), Fig S3F (C). Red rectangular indicates the parts of the blots that are used in figures.

**S1 File. Supplemental Table 1** Scoring data of Centrin-3 foci vs. CEP152, CEP170 and γ-tubulin foci of -dox, 24H PLK4 induced and 48H PLK4 induced cells in Fig. S1D, Fig. 1E and Fig. 1F.

**S2 File. Supplemental Table 2** Scoring data of planar vs. circular oriented CRCs presented in Figs. S2E, S2F, S2G, 2G and 2H.

**S3 File. Supplemental Table 3** Scoring data of CR and CRCs distances in Figs. S3C, S3E, 3B, 3C, 3D and 3E.

**S4 File. Supplemental Table 4** Scoring data of Centrin-3 foci vs. CEP152 foci in Fig. 4C.

## Acknowledgements

We are thankful to Sercin Karahuseyinoglu and KUTTAM-CMIC team for their technical assistance on high resolution imaging. The authors gratefully acknowledge the use of the services and facilities of the Koç University Research Center for Translational Medicine (KUTTAM), funded by the Presidency of Turkey, Presidency of Strategy and Budget.

## Author Contributions

Conceptualization: SCO and CAA, Investigation: SCO, BMK, EC and AAC, Methodology: SCO and BMK, Visualization: SCO, Project Administration: CAA, Funding Acquisition: CAA and SCO, Writing - original draft: SCO, Writing - review & editing: CAA.

## Notes

### Competing Interest Statement

The authors have declared no competing interest.

