## Supplementary figures and images for "Prolonged over-expression of PLK4 amplifies centrosomes through formation of inter-connected centrosome rosette clusters"

### Supplemental Figure 1

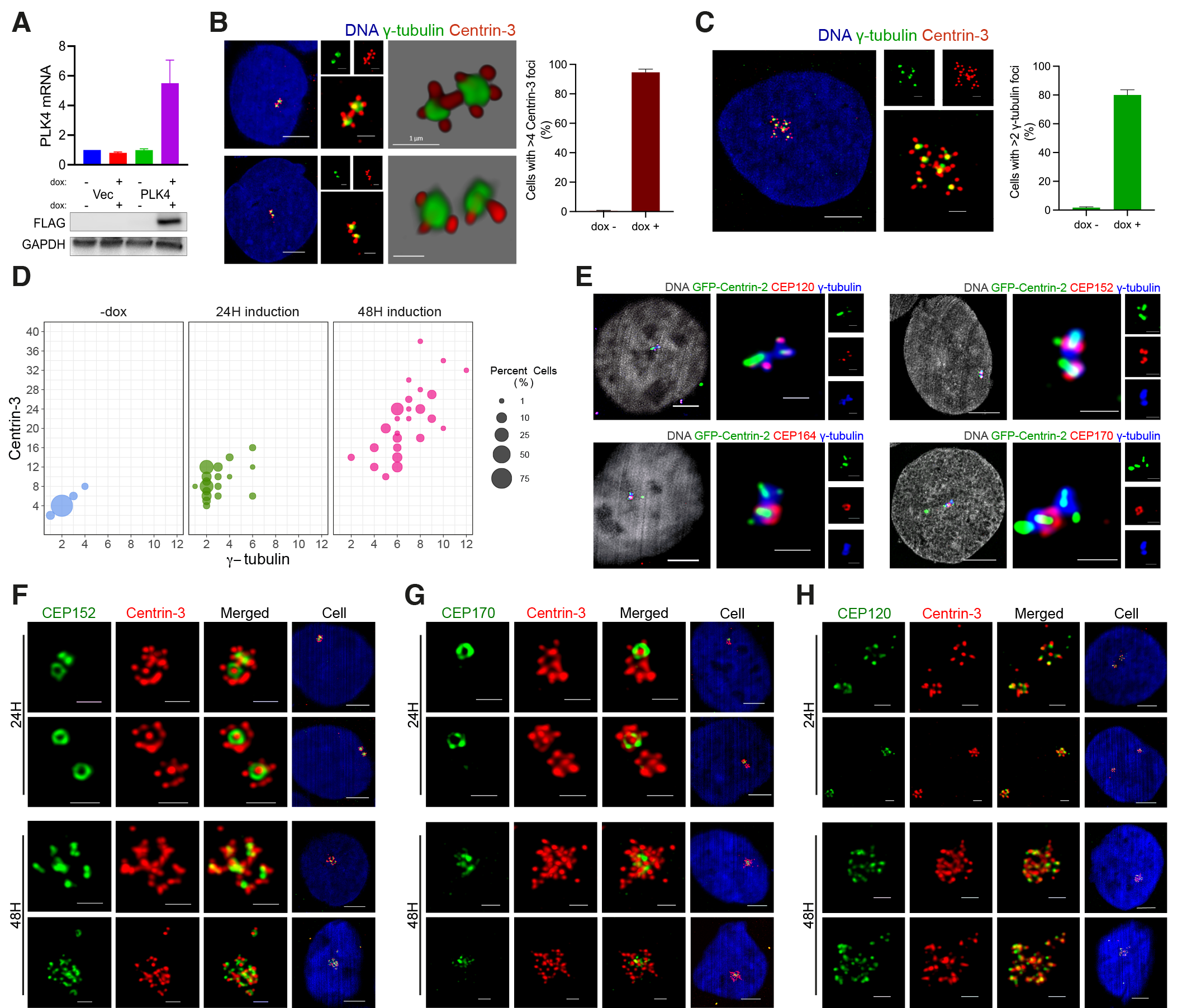

### Supplemental Figure 2

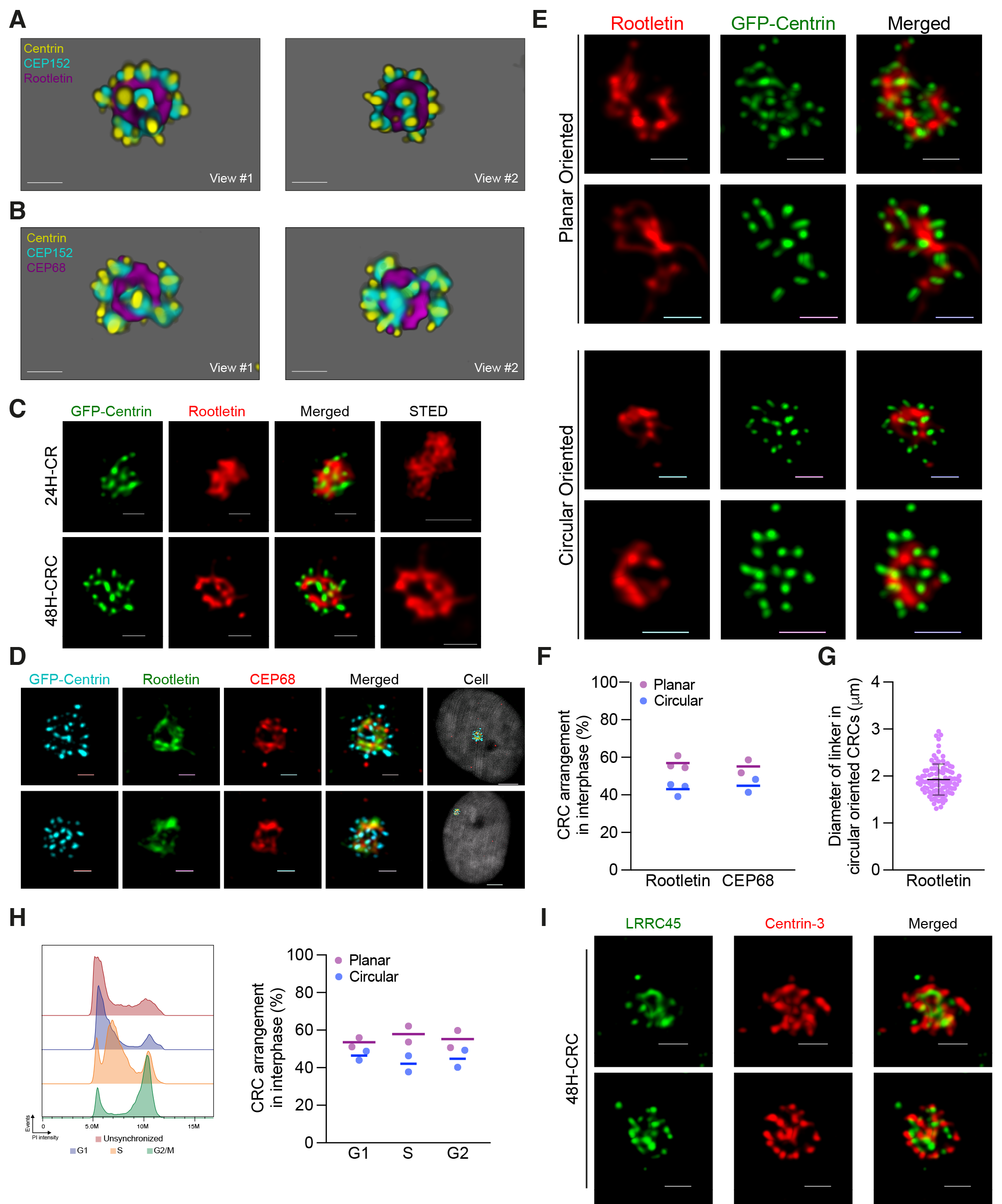

### Supplemental Figure 3

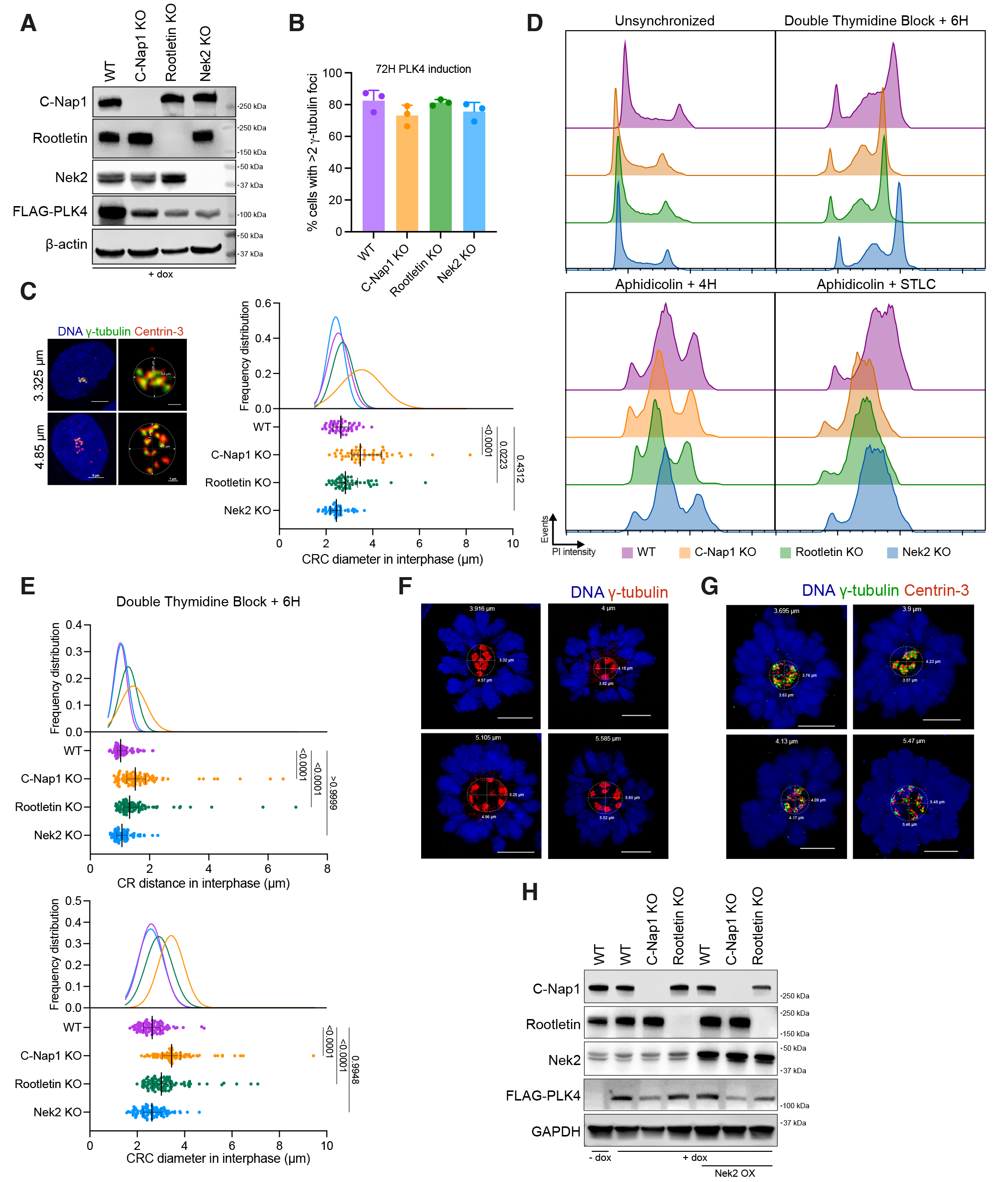

### Supplemental Figure 4

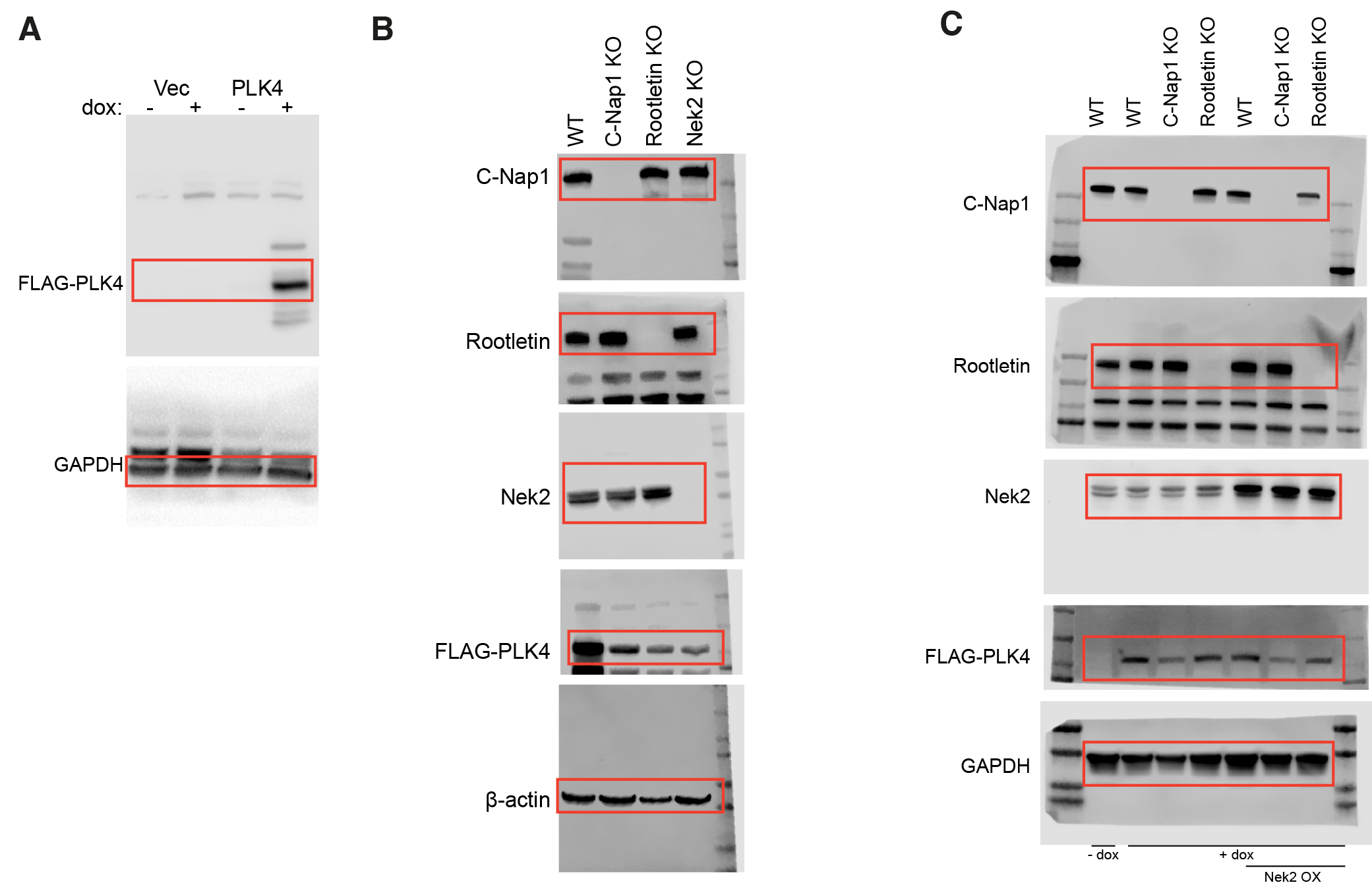
